# Co-occurrence or dependence? Using spatial analyses to explore the interaction between palms and triatomines (Chagas disease insect vectors)

**DOI:** 10.1101/644385

**Authors:** Johan M. Calderón, Camila González

**Affiliations:** Centro de Investigaciones en Microbiología y Parasitología Tropical (CIMPAT), Departamento de Ciencias Biológicas, Universidad de Los Andes, Bogotá D.C., Colombia

## Abstract

**Background:** Triatomine kissing bugs are responsible for the vectorial transmission of the parasite *Trypanosoma cruzi,* etiological agent of Chagas disease, a zoonosis affecting 10 million people and with 25 million at risk of infection. Triatomines are associated with particular habitats that offer shelter and food. Several triatomine species of the *Rhodnius* genus have close association with palm crowns, where bugs can obtain blood from the associated fauna. The *Rhodnius* - palm interaction has been reported in several places of Central and South America. However, the association in the distributions of *Rhodnius* species and palms has not been quantitatively determined.

**Methodology/Principal Findings:** Broad distributions of eight *Rhodnius* species and 16 palm species with *Rhodnius*-infestation reports were estimated using Ecological Niche Models. *Rhodnius* species distributions in their total range were compared to their distributions in areas with palms. *Rhodnius* species presence was found to be higher in areas with palms. However, that tendency notoriously depended on palm species. *Rhodnius* species presence increased several times in areas with particular palm species. Moreover, a possible relationship was found between *Rhodnius* and palm species richness, indicating the Amazon region as the convergent region where several *Rhodnius* and palm species intersected. Finally, palm distribution was evaluated as predictor of *Rhodnius* species distributions, but their inclusion in the distributions models did not improve their performance.

**Conclusions/Significance:** The distributions of some *Rhodnius* and palm species showed a high spatial association, which can be based on species interaction or niche similarity. Based on distribution convergence, the Amazon region appear to be the origin of the *Rhodniu*s-palm association. The direct relationship between palms and *Rhodnius* species richness could be based on the habitat heterogeneity offered by different palm species. Despite spatial association, palm presence would not be a relevant predictor of *Rhodnius* species distributions in comparison to other environmental variables. Inclusion of other input data as hosts’ distribution could help to increase model predictability.

**Author summary:** The infestation of palms with *Rhodnius* genus kissing bugs (Chagas disease vectors) is important from the public health perspective, since insects living in palms can infest nearby houses. The migration of these bugs to households could threaten vector control programs since reinfestation of treated dwellings can occur. Association between *Rhodnius* and palms species distributions has been previously suggested but never quantitatively determined. The strong association between one palm species and one *Rhodnius* species can be used as a factor to predict the presence of *Rhodnius* bugs in definite areas. In this study, we estimated by models the distributions of eight *Rhodnius* species and 18 *Rhodnius*-infested palm species. *Rhodnius* distributions models showed a biased presence toward areas with certain palm species. That specific association was very strong in some cases; however, the presence of associated palm species was used in *Rhodnius* distributions models, but that did not improve the predictability of the models. Palm presence appear to be not essential for the *Rhodnius* current distribution because they could inhabit other habitats; but that association could be relevant to the *Rhodnius* evolutionary and biogeographic history.

## Introduction

Triatomine kissing bugs are responsible for the vectorial transmission of the parasite *Trypanosoma cruzi,* etiological agent of Chagas disease, a zoonosis affecting 10 million people and with 25 million at risk of infection [1]. Triatomines show associations with particular habitats that offer shelter and food [2]; this association can be specific to one type of habitat as occurs with *Psammolestes* triatomines living in bird nests, or to several types as *Triatoma sordida* which can be found in rock piles, hollow trees, and human dwellings [3]. Several species belonging to the genus *Rhodnius,* for instance, have been found in close association with palms in its sylvatic cycle [4]. Palm crowns have been suggested as suitable places for an associated-fauna where *Rhodnius* can obtain blood, and *Rhodnius neglectus* and *Rhodnius nasutus*, for example, have been reported to feed from birds [5,6] using palms as nesting sites. Furthermore *Didelphis marsupialis* one of the most competent hosts for *T. cruzi* eats palm-tree fruits and rests in the clefts between palm stipe and fronds [7].

The fact that palms are infested with Chagas disease vectors is important from the public health perspective, since insects living in palms can infest nearby houses [8,9]. Triatomines are also capable of colonizing non-native palm species, as the ones in plantations or used for garden decoration, increasing the risk of domiciliation of the disease [10,11]. In addition, the use of palms in households (e.g. dry leaves for roof thatching) could have a major role in the domiciliation of the disease [4,12]. The migration of kissing bugs to households could threaten vector control programs conducted as a complement during Chagas disease control initiatives, since reinfestation of treated dwellings can occur [13].

*Rhodnius* species are distributed from Central America to northern Argentina, being the Amazon region the zone with the highest number of species [14]. They are primarily associated with palms, but also occur in bird nests, mammal burrows, peridomestic and domestic habitats [15]. Even one species, *Rhodnius domesticus,* has been reported in bromeliads and hollow trees, but not in palms [4]. *Rhodnius* species distribution shows a segregated pattern across America: *Rhodnius prolixus*, *Rhodnius pallescens* and *Rhodnius neivai* are found in Central America and northern South America; *Rhodnius pictipes*, *Rhodnius robustus* and *Rhodnius brethesi* in the Amazon region; *Rhodnius nasutus* and *Rhodnius neglectus* in northeastern and central Brazil; *Rhodnius ecuadoriensis* in Ecuador and northern Perú; and *Rhodnius stali* in Bolivia [14,16,17].

Palms are mainly distributed in the tropics, but a few species reach subtropical zones both in the northern and southern hemisphere [18]. The highest palm concentration is found in the intertropical zone, being Asian and American tropics the richest areas in terms of species number. In America, palms are distributed from southern United States to northern Argentina and central Chile [18]. From 550 palm species naturally occurring in America [19], 22 have been reported infested by *Rhodnius* triatomines [20], and the genus *Attalea*, itself has five species reported as infested: *At. butyracea*, *At. maripa*, *At. oleifera*, *At. phalerata* and *At. speciosa*. *Attalea butyracea*, is a species extensively studied as an ecotope for *Rhodnius* triatomines and for Chagas disease transmission [13,21–23].

The association *Rhodnius* - palm has been observed and reported in several places of Central and South America [20]. This has led to the conclusion, as in Gaunt & Miles in 2000 [2], that sylvatic *Rhodnius* distribution should broadly coincide with palms distribution. However, this coincidence has not been explicitly evaluated. *Rhodnius* and palms distributions have not been compared yet by a quantitative evaluation to determine if the geographical presence of both organisms is the result of choice or chance.

Both *Rhodnius* and palms spatial distributions have been depicted using outline maps to define the limits of the distributions [16,18,19,24], or by locality reports, summarizing the places where the species have been found [25–29]. Outline maps vary in accuracy based on how well known the distribution is and how precisely the author incorporated the information. Locality reports are accurate, but they can only show a fraction of the area where the species live [30]. With both sources of information, it is difficult to make a quantitative comparison covering the complete area of distribution. One very used alternative is to estimate *Rhodnius* and palm distributions through modeling. Distribution models extrapolate information of species location records in space and time, usually based on statistical models [31]. One type of these models are the Ecological Niche Models (ENM); they allow to predict species presence in a region based on habitat suitability [31]. Locations with similar environmental conditions to those where the species was observed are considered as suitable habitats for the species presence.

ENM have been previously used to estimate *Rhodnius* species distribution: *R. neglectus*, *R. nasutus*, *R. pictipes* and *R. robustus* in Brazil [32–35]; *R. pallescens* in Colombia, Panamá, Costa Rica, and Nicaragua [36,37], and *Rhodnius prolixus* in Colombia [37] and Guatemala [38]. Those studies did not considered biological interactions as predictors for *Rhodnius* distributions. Based on the available information and *Rhodnius* ecology, palm presence could increase the predictability of *Rhodnius* ENM adding information about possible ecotopes. Including biological interactions have been used in previous studies improving the ENM predictability [39,40], and showing, in some cases, a higher effect from the biotic predictors compared to the abiotic [41]. Species interactions can be included in models by limiting the predicted distribution of one species to the distribution of another [42] or by including the presence of one species as a predictor [41,43–45].

The aim of this study was to quantitatively assess the association between *Rhodnius* and palms distributions using ENM. First, to determine if *Rhodnius* species presence is biased toward areas with palms, its presence was compared in the entire modeled area and in zones with palms. Second, to identify if *Rhodnius*-palm association is species-specific, prevalence comparisons were carried out discriminating by palm species. And third, to determine if palm presence could be considered as predictor of *Rhodnius* species distributions, *Rhodnius* ENM were run again with palm distributions as environmental predictors. The performance of ENMs with or without palm distributions were compared to identify any prediction improvement caused by the addition of biological interactions.

## Methods

### Ecological niche models

To estimate *Rhodnius* and palms potential distributions, ENM were carried out for eight *Rhodnius* species collected in palms, and the 16 palm species where those kissing bugs were found (Table 1). Four additional *Rhodnius* species (*R. barretti*, *R. brethesi*, *R. neivai*, and *R. stali*), and one palm species (*Copernicia tectorum)* were initially included in the study but due to the low number of occurrences (less than 19 occurrences), their spatial analyses were not performed.

**Table 1.**
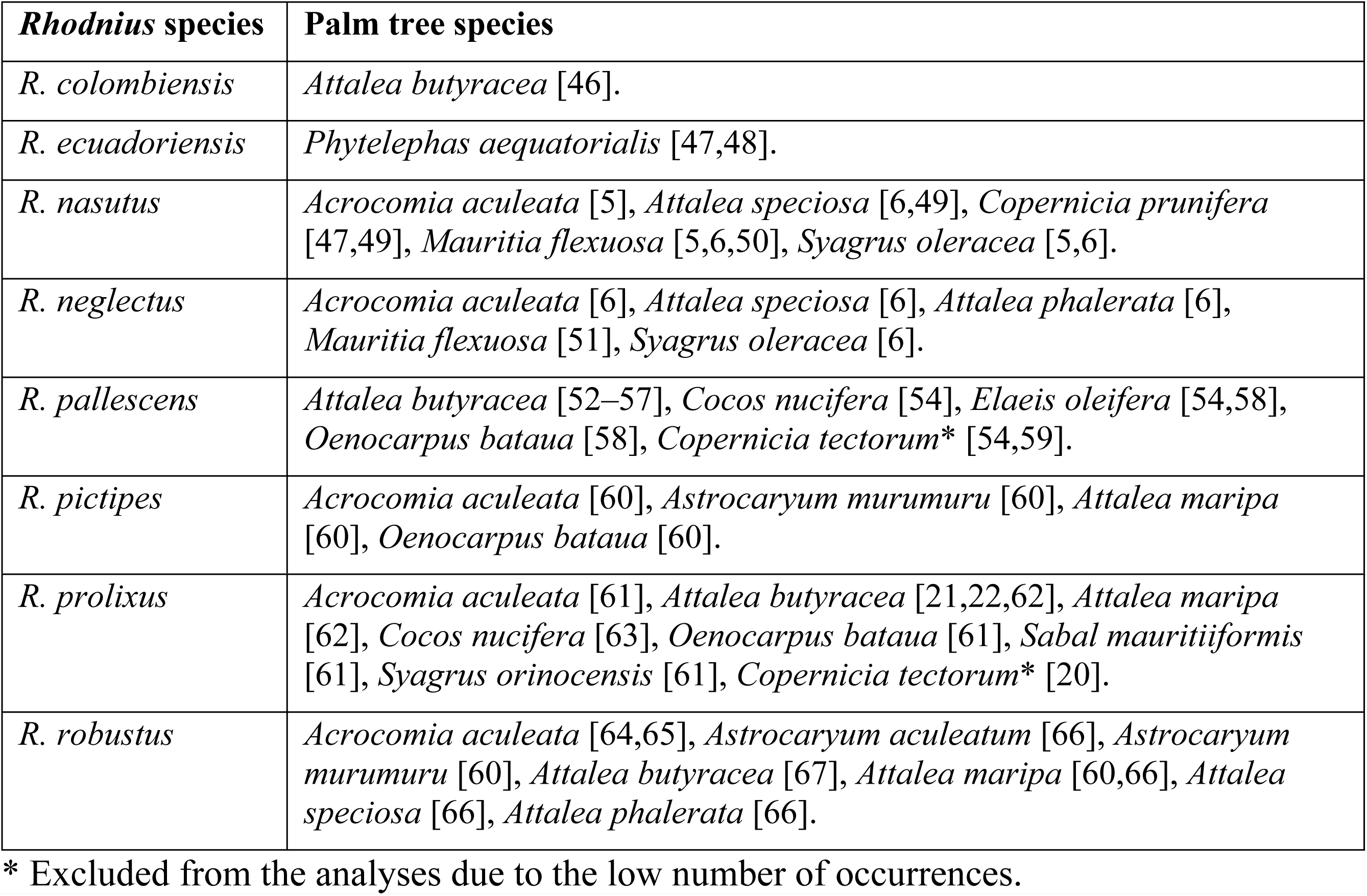
*Rhodnius* species found infesting palm trees.

| *Rhodnius* species | Palm tree species |
| --- | --- |
| *R. colombiensis* | *Attalea butyracea* [46]. |
| *R. ecuadoriensis* | *Phytelephas aequatorialis* [47,48]. |
| *R. nasutus* | *Acrocomia aculeata* [5], *Attalea speciosa* [6,49], *Copernicia prunifera* [47,49], *Mauritia flexuosa* [5,6,50], *Syagrus oleracea* [5,6]. |
| *R. neglectus* | *Acrocomia aculeata* [6], *Attalea speciosa* [6], *Attalea phalerata* [6], *Mauritia flexuosa* [51], *Syagrus oleracea* [6]. |
| *R. pallescens* | *Attalea butyracea* [52–57], *Cocos nucifera* [54], *Elaeis oleifera* [54,58], *Oenocarpus bataua* [58], *Copernicia tectorum* * [54,59]. |
| *R. pictipes* | *Acrocomia aculeata* [60], *Astrocaryum murumuru* [60], *Attalea maripa* [60], *Oenocarpus bataua* [60]. |
| *R. prolixus* | *Acrocomia aculeata* [61], *Attalea butyracea* [21,22,62], *Attalea maripa* [62], *Cocos nucifera* [63], *Oenocarpus bataua* [61], *Sabal mauritiiformis* [61], *Syagrus orinocensis* [61], *Copernicia tectorum* * [20]. |
| *R. robustus* | *Acrocomia aculeata* [64,65], *Astrocaryum aculeatum* [66], *Astrocaryum murumuru* [60], *Attalea butyracea* [67], *Attalea maripa* [60,66], *Attalea speciosa* [66], *Attalea phalerata* [66]. |

\* Excluded from the analyses due to the low number of occurrences.

#### Occurrences

*Rhodnius* occurrences (i.e. locations were the species were found) were obtained from “DataTri”, a database of American triatomine species occurrences [68]. Palm tree occurrences were obtained from the Global Biodiversity Information Facility (GBIF; downloaded in October 2018) using the “gbif” function of the “dismo” R package [69]. Occurrences with both geographical coordinates were selected, and all the duplicated records were removed.

A depuration of the compiled database was performed, and *Rhodnius* and palm tree occurrences were checked to correspond with previous geographical distributions reported in the literature [18,19,72,73,25,27,28,33,36,46,70,71]. Occurrences in altitudes outside species limits were omitted (maximum altitude above sea level: *R. ecuadoriensis* 1500 m, *R. nasutus* 700 m, *R. neglectus* 800 m, *R. pallescens* 400 m, *R. pictipes* 1100 m, *R. prolixus* 2000 m, *R. robustus* 1200 m [16], *R. colombiensis,* value not found. Palm species: *Ac. aculeata* 1300 m, *As. murumuru* 900 m, *At. butyracea* 1000 m, *At. maripa* 600 m, *At. phalerata* 1000 m, *Cc. nucifera* 1800 m, *E. oleifera* 300 m, *M. flexuosa* 900 m, *O. bataua* 1000 m, *P. aequatorialis* 1500 m, *Sa. mauritiiformis* 1000 m, *Sy. oleracea* 800 m, and *Sy. orinocensis* 400 m [18,19,73], for *As. aculeatum, At. speciosa,* and *Cp. prunifera*, the value was not found). *Rhodnius prolixus* occurrences in Central America were excluded from the study since the species was only related to domestic transmission; they are no longer found in previous reported areas as a possible consequence of vector control initiatives [74], and the species presence has not been associated with palm trees [14,17,75].

To reduce the effect of sampling bias in the occurrence data set, spatial thinning was performed with the “spThin” R package [76] using a minimum nearest neighbor distance greater than or equal to 10 km.

#### Environmental variables

The set of environmental variables was composed by the 19 bioclimatic variables from WorldClim [77], three topographic variables (slope, aspect, and topographic position index (TPI); calculated from the GTOPO30 DEM [78]), and 42 variables with remote sensing information of land surface temperature (LST), normalized difference vegetation index (NDVI), and middle infrared radiation (MIR). The remote sensing variables were calculated from AVHRR (Advanced Very High-Resolution Radiometer) images and processed by the TALA group (Oxford University, UK) using the temporal decomposition of Fourier [79]. Pearson correlation coefficient was calculated among variables to avoid collinearity, and from a group of variables showing high correlation (i.e. r absolute value bigger than 0.7), only one variable was selected. This selection was based on which variable grouped more temporal information (e. g. yearly over monthly). The 12 selected environmental variables included six bioclimatic variables (1, 2, 12, 15, 18), three topographic variables (slope, aspect, TPI), and five remote sensing variables (Mean LST, LST annual phase, mean NDVI, and NDVI variance). Correlation was double-checked by the Variable Inflation Factor (VIF), obtaining values lower than three for every variable. The spatial resolution of all the layers was 2.5° (approximately 5Km^2^).

#### Modeling and evaluation

Pseudo-absences for the *Rhodnius* models were obtained from the occurrences of all the triatomines species but the modeled one. Those pseudoabsences give a higher discriminative ability than background data because the records are concentrated into the accessible area for triatomines. The geographical extent used for each *Rhodnius* model was the species range reported in the literature [25,27,28,33,36,46,70–72]. As modeling algorithms, five techniques were used: Generalized Linear Models (GLM), Generalized Boosting Models (GBM), Generalized Additive Models (GAM), Maximum Entropy (MaxEnt), and Random Forest (RF). Modeling was carried out with “Biomod2” R package [80]; this package allows to run and evaluate several algorithms in parallel. Default options were chosen for each algorithm except for MaxEnt. Variation in the regularization multiplier (β) and feature classes in MaxEnt have shown to affect significantly the model performance [81]. Several β values (0.02, 0.1, 0.46, 1, 2.2, and 4.6) and feature classes (linear, quadratic and product) were tested for each species with the “ENMeval” R package [82], and the options giving the lowest Akaike Information Criterion (AIC) were selected.

Considering palm tree species, all models were carried out in the same calibration area, from Nicaragua to northern Argentina. This area includes the distribution of all eight *Rhodnius* species evaluated. In contrast to *Rhodnius* models, background data was used (10,000 random points) eliminating the points coinciding with palm presence.

For each modeled species, ENM were run ten times with different presence and pseudo-absence subsamples to test robustness [83]. Each time, 80% of the occurrences and pseudo-absences were randomly chosen for training the model and the left 20% of the occurrences used for testing. Model evaluation was based on two methods, partial area under the ROC curve (pAUC) [84] and omission rates. The first one was calculated with two omission levels, 0.10 and 0.50, later obtaining the ratio between both pAUCs. The process was repeated 100 times (using bootstrap subsampling) to estimate 95% confidence intervals. Ten-percentile and zero-percentile training omission rates (proportion of testing occurrences omitted with each threshold) were calculated along with the presence prevalence (proportion of presence area compared to the entire modeled area).

Final outputs used for comparing *Rhodnius* and palm distributions were obtained with 100% of the occurrences (and pseudo-absences for *Rhodnius* models). The entire occurrence set gives all the available information to the model for being as accurate as possible. Binary maps were obtained from these outputs using the 10-percentile threshold. To assemble the predictions given by the five algorithms, binary maps were summed, and presence was defined as the resulting area where three or more algorithms coincided. This procedure could be conservative, letting some presence records out, but it allowed to work with predicted presences with high amount of support.

### Association between *Rhodnius* species and palm trees distributions

To determine if *Rhodnius* species presence is biased toward areas where palms are present, both estimated distributions were compared using prevalence. Species prevalence is the proportion of species presence in a definite area. Prevalence for each *Rhodnius* species was calculated in the total area and in the areas with predicted palm presence. Both prevalence values were compared using odds ratio and calculating 95% confidence intervals. Whether odds ratio was bigger than one, prevalence in areas with palms was higher than in the entire area. Higher the odd ratio values, higher the possible *Rhodnius* - palms association. Palm presence in the models was defined as the presence of at least one palm species, regardless of the species.

Then, to identify whether *Rhodnius*-palm association depend on palm species, *Rhodnius* species prevalence was calculated in areas with palm presence discriminating by palm species. Obtained values were compared with the total prevalence again using the odds ratio and calculating 95% confidence intervals. Since *Rhodnius* prevalence in palm areas can be affected by palm prevalence, palm species with prevalence values smaller than 0.10 were excluded of the analyses. To compare *Rhodnius* and palm distributions, they must be in the same spatial extension, so all the palm distributions were cropped to the extension of each *Rhodnius* species model using the function “crop” of the R package “raster” [85]. Using the *Rhodnius* species extension avoided to include overpredicted areas and limit the comparison to the geographical range reported in the literature.

### *Rhodnius* models with palm trees distributions as predictors

To determine if palm presence could be considered as a predictor of *Rhodnius* species presence, *Rhodnius* models were run again including the palm tree predicted presence as a layer. Binary palm distributions used for each *Rhodnius* species were those of palm species showing a high association (odd ratio values higher than 2). The modeling process and evaluation methods were the same as described for the previous models. Evaluation statistics were compared between *Rhodnius* models with and without palm trees distributions as predictors.

## Results

### Ecological niche models

In *Rhodnius* species, the number of occurrences varied from 19 in *R. colombiensis* to 352 in *R. prolixus* (Table 2). Spatial distribution also showed great variation: *Rhodnius robustus* and *R. pictipes* occurrences encompassed the widest area including several countries (more than 6,500,000 km^2^); while *R. ecuadoriensis* and *R. colombiensis* presence occupied much smaller areas (less than 50,000 km^2^) (Fig. 1). Considering model performance, all the *Rhodnius* models had pAUC ratios significantly higher than the null model line (i.e. omission 0.50) showing a good ability to discriminate (Table 2). Omission rates had a contrasting performance: ten percentile omission rates were higher than the expected value in all the models, and in some species as *R. nasutus*, *R. colombiensis* and *R. ecuadoriensis* values were exceptionally high. Zero percent omission rates were different, being very close to the expected values in all the species except for *R. colombiensis* and *R. ecuadoriensis*. Three bioclimatic variables: annual mean temperature (Bio 1), annual precipitation (Bio 12), and precipitation seasonality (Bio15), and one remote sensing variable, NDVI variance, were the most influential variables in five of the eight *Rhodnius* species. In contrast, topographic variables as aspect, slope and TPI showed low influence in the models.

**Fig 1.**
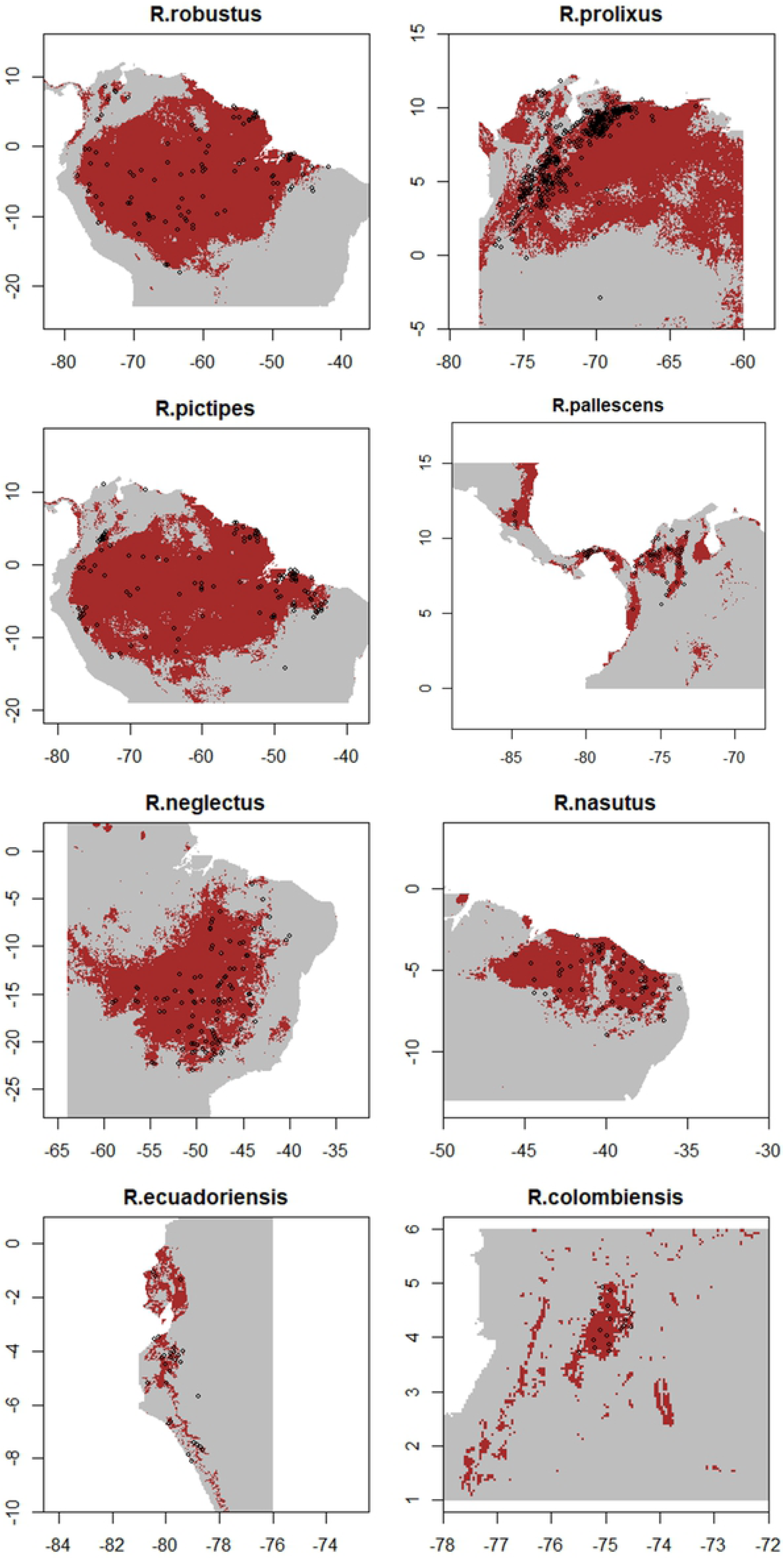
*Rhodnius* species predicted distribution. Brown: Predicted presence. Gray: Predicted absence. Black points: Species occurrences. Presences were predicted in at least three algorithms based on the 10% omission rate threshold. Horizontal axis: Longitude, Vertical axis: Latitude. Maps constructed with the raster R package [85].

**Table 2.** Performance statistics in *Rhodnius* ENM.

**A. With environmental variables**
| Species | Occurr | Partial AUC ratio |  | Omission rate |  |  |  |
| --- | --- | --- | --- | --- | --- | --- | --- |
|  |  | Median* | 95%CI | 10%<sup>1</sup> | Prev 10% | 0%<sup>1</sup> | Prev 0% |
| *R. robustus* | 96 | 1.366 | 1.148-1.530 | 0.1500 | 0.5048 | 0.0250 | 0.5862 |
| *R. prolixus* | 352 | 1.474 | 1.313-1.606 | 0.1831 | 0.5147 | 0.0070 | 0.9002 |
| *R. pictipes* | 117 | 1.563 | 1.351-1.733 | 0.1875 | 0.5345 | 0 | 0.7130 |
| *R. pallescens* | 67 | 1.668 | 1.296-1.859 | 0.1623 | 0.1623 | 0.0360 | 0.5124 |
| *R. neglectus* | 101 | 1.626 | 1.458-1.765 | 0.1667 | 0.2705 | 0 | 0.5979 |
| *R. nasutus* | 61 | 1.576 | 1.332-1.750 | 0.5000 | 0.1991 | 0 | 0.3673 |
| *R. ecuadoriensis* | 28 | 1.728 | 1.549-1.879 | 0.5000 | 0.1421 | 0.2500 | 0.2185 |
| *R. colombiensis* | 19 | 1.934 | 1.439-1.983 | 0.3750 | 0.0267 | 0.1250 | 0.0377 |

**B. With environmental variables and palm distributions**
| Species | Partial AUC ratio |  | Omission rate |  |  |  | Importance of palm distributions<sup>2</sup> |
| --- | --- | --- | --- | --- | --- | --- | --- |
|  | Median* | 95%CI | 10%<sup>1</sup> | Prev 10% | 0%<sup>1</sup> | Prev 0% |  |
| *R. robustus* | 1.358 | 1.210-1.506 | 0.2000 | 0.4640 | 0.0250 | 0.5939 | 13 |
| *R. prolixus* | 1.540 | 1.491-1.645 | 0.1796 | 0.4538 | 0.0100 | 0.8847 | 5 |
| *R. pictipes* | 1.282 | 1.007-1.494 | 0.2292 | 0.5641 | 0.0312 | 0.7054 | 6 |
| *R. pallescens* | 1.686 | 1.338-1.880 | 0.2679 | 0.1648 | 0.0536 | 0.5539 | 3 |
| *R. neglectus* | 1.504 | 0.9542-1.702 | 0.2857 | 0.2726 | 0.0357 | 0.6307 | 5 |
| *R. nasutus* | 1.638 | 1.363-1.794 | 0.4423 | 0.2355 | 0.0769 | 0.4174 | 13 |
| *R. ecuadoriensis* | 1.749 | 1.467-1.933 | 0.3750 | 0.1657 | 0.1667 | 0.2185 | 13 |
| *R. colombiensis* | 1.922 | 0.9954-1.999 | 0.3750 | 0.0222 | 0.3125 | 0.0258 | 3 |

<sup>1</sup> Median value for the five used algorithms. With each algorithm, value was the median of the ten repetitions carried out with different training and testing data set.
<sup>2</sup> Median value for the importance variable ranks with all the algorithms.

Most of the *Rhodnius* models predicted an area of distribution adjusted to the occurrence points (Fig 1). However, *R. prolixus*, *R. pallescens*, and *R. colombiensis*, models showed over-prediction areas outside occurrences (Fig 1). For instance, *R. prolixus* model predicted presence into Venezuelan and Colombian Amazon, *R. pallescens* model in the eastern Nicaragua, and *R. colombiensis* model in several zones of the Cauca river valley, where the species have not been found [36,46,75].

In palm species, the number of occurrences varied from 24 in *P. aequatorialis* to 326 in *O. bataua* (Table 3). Spatial distribution showed a high variation among species: *C. nucifera* and *M. flexuosa* had very wide distributions comprehending almost half of the entire modeled area (presence in more than 6,500,000 km^2^); meanwhile, species as *P. aequatorialis* and *Sa. mauritiiformis* had narrow distributions covering only small definite areas (less than 140.000 km^2^) (Fig. 2).

**Fig 2.**
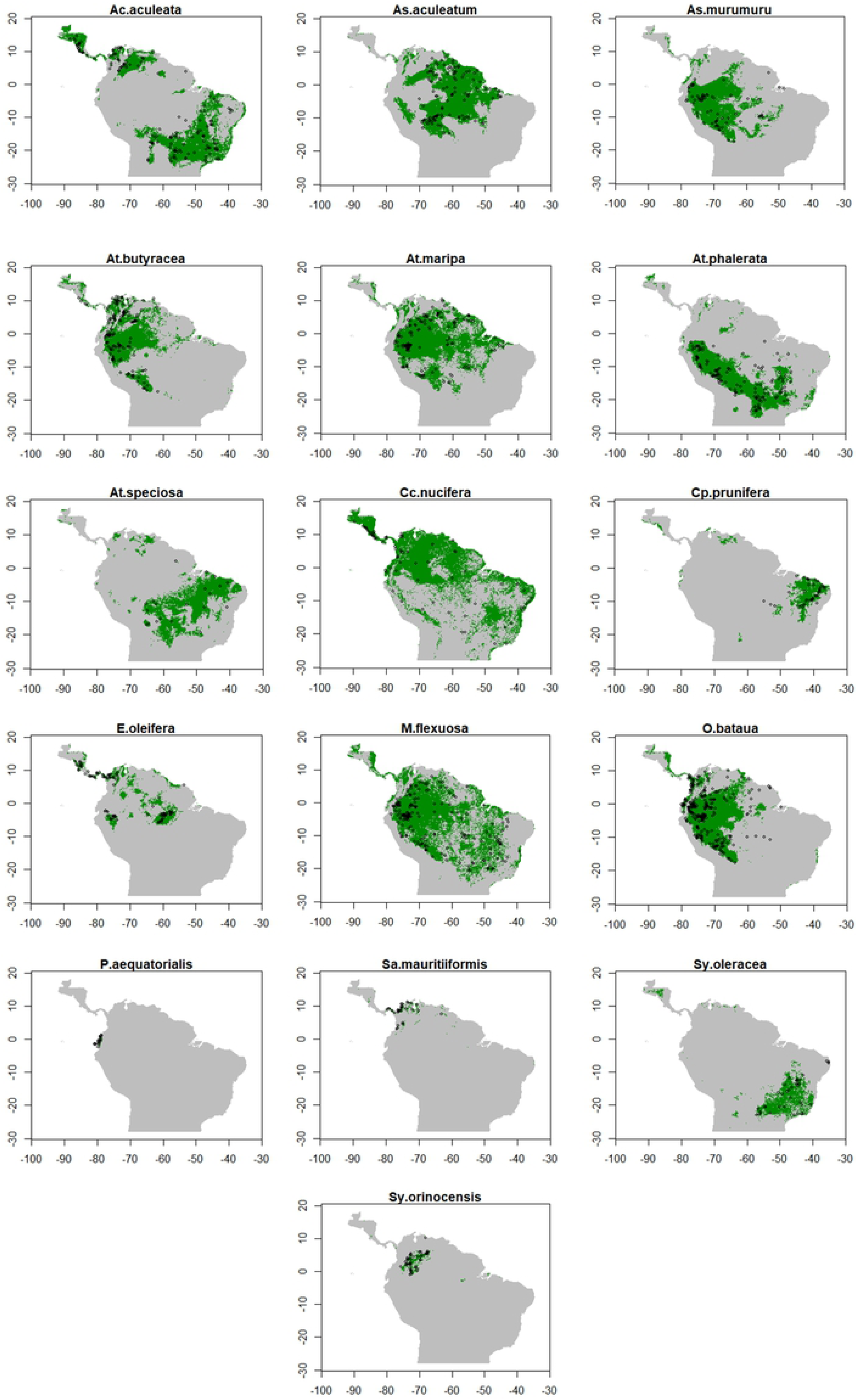
Palm species predicted distribution. Green: Predicted presence. Gray: Predicted absence. Black points: Occurrences. Presence was predicted in at least three algorithms based on the 10% training omission threshold. Horizontal axis: Longitude, Vertical axis: Latitude. Maps constructed with the raster R package [85].

**Table 3.** Performance statistics in palms ENM.

| Species | Occur. | Partial AUC ratio |  | Omission rate |  |  |  |
| --- | --- | --- | --- | --- | --- | --- | --- |
|  |  | Median* | 95%CI | 10%* | Prev 10% | 0%* | Prev 0% |
| *Ac. aculeata* | 154 | 1.676 | 1.510-1.793 | 0.1613 | 0.2795 | 0 | 0.6601 |
| *As. aculeatum* | 119 | 1.641 | 1.467-1.774 | 0.2083 | 0.2931 | 0.0417 | 0.4230 |
| *As. murumuru* | 75 | 1.689 | 1.399-1.858 | 0.2333 | 0.2299 | 0.0667 | 0.3673 |
| *At. butyracea* | 153 | 1.765 | 1.644-1.869 | 0.1935 | 0.2145 | 0 | 0.4183 |
| *At. maripa* | 159 | 1.672 | 1.493-1.800 | 0.1875 | 0.3240 | 0 | 0.5232 |
| *At. phalerata* | 150 | 1.710 | 1.535-1.846 | 0.1833 | 0.2425 | 0 | 0.6741 |
| *At. speciosa* | 20 | 1.595 | 0.9517-1.999 | 0.4375 | 0.3189 | 0 | 0.4624 |
| *Cc. nucifera* | 74 | 1.499 | 1.145-1.783 | 0.1667 | 0.4830 | 0 | 0.7177 |
| *Cp. prunifera* | 45 | 1.922 | 1.759-1.981 | 0.1111 | 0.0696 | 0 | 0.1363 |
| *E. oleifera* | 93 | 1.868 | 1.704-1.951 | 0.2632 | 0.0647 | 0 | 0.2226 |
| *M. flexuosa* | 206 | 1.576 | 1.406-1.726 | 0.1667 | 0.4368 | 0 | 0.7357 |
| *O. bataua* | 323 | 1.781 | 1.699-1.851 | 0.1154 | 0.2272 | 0 | 0.5771 |
| *P. aequatorialis* | 24 | 1.994 | 1.497-1.999 | 0.8000 | 0.0011 | 0.4000 | 0.0014 |
| *Sa. mauritiiformis* | 30 | 1.769 | 1.163-1.990 | 0.5833 | 0.0099 | 0.2500 | 0.0292 |
| *Sy. oleracea* | 38 | 1.762 | 1.451-1.929 | 0.2500 | 0.1090 | 0.1875 | 0.2082 |
| *Sy. orinocensis* | 48 | 1.966 | 1.829-1.984 | 0.2000 | 0.0248 | 0 | 0.0766 |

\* Median value for the five used algorithms. With each algorithm, value was the median of the ten repetitions carried out with different training and testing data set.

Considering performance, all the palm models showed pAUC ratios significantly higher than the null model line except in *A. speciosa* (Table 3). Like *Rhodnius* models, 10 percentile omission rates were higher than the expected value, but the zero percentile omission rate were very close to the expected one. Three species showed very high omission rates values with both thresholds: *P. aequatorialis*, *Sa. mauritiiformis* and *Sy. oleracea*. *Attalea speciosa*, showed a very high 10 percentile omission rate but low zero percentile. Every palm model predicted an area of distribution adjusted to the occurrence points (Fig 2), and no clear over-prediction was identified in any model. Considering predictors for palm species distributions, annual mean temperature (Bio 1) was an important variable in eleven palm species, and three variables, annual precipitation (Bio 12), precipitation seasonality (Bio 15), and precipitation of the warmest quarter (Bio 18), were important in eight palm species. In contrast, topographic variables showed low influence in the models.

As an alternative to decrease the ten-percentile omission rates, ENMs were repeated using as layers, the first 16 PCAs obtained from the original 42 variables (which covered 90% of the environmental variation). However, omission rates did not improve (S1 Table) and the initial ENM were used for the further analysis.

### Association between *Rhodnius* species and palms distributions

*Rhodnius* species prevalence was higher in areas with palm presence compared to the entire area, except for two species, *R. prolixus* and *R. colombiensis* (Table 4). However, differences between prevalence values were small; all the odds ratios were close to 1. Palm prevalence (presence of at least one palm species) was very high in all the *Rhodnius* species distribution areas (Fig 3). In some cases, as in *R. robustus* and *R. pallescens*, presence of palm trees covered almost the entire area (Table 4). In *R. ecuadoriensis*, with the smallest palm prevalence, palm presence was wide distributed comprehending more than a half of the total area.

**Fig 3.**
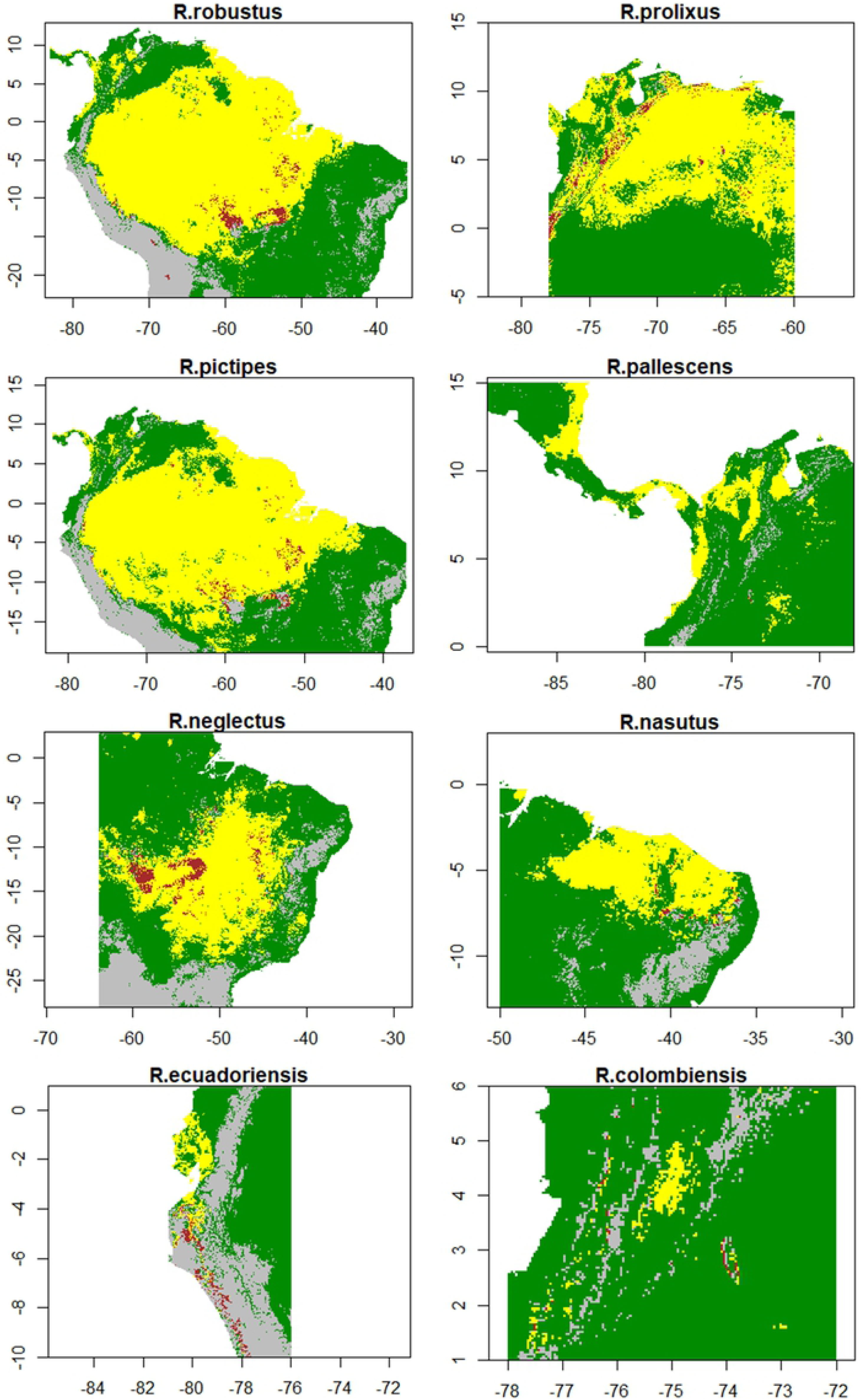
*Rhodnius* and palm trees distributions. Yellow: Presence of both the Rhodnius species and palms. Brown: Only the Rhodnius species. Green: Only palms. Gray: Both absences. Palms presence corresponded to the presence of at least one palm species. All shown presences were predicted in at least three algorithms based on the 10% training omission threshold. Horizontal axis: Longitude, Vertical axis: Latitude. Maps constructed with the raster R package [85].

**Table 4.** Rhodnius species prevalence in the entire distribution area and in palm trees areas.

| *Rhodnius* species | Species prevalence | Palm prevalence | Species prevalence in palm areas | Odds ratio 95% CI* |
| --- | --- | --- | --- | --- |
| *R. robustus* | 0.5076 | 0.8658 | 0.5701 | 1.277-1.296 |
| *R. prolixus* | 0.4772 | 0.9777 | 0.4726 | 0.9678-0.9955 |
| *R. pictipes* | 0.5306 | 0.8867 | 0.5875 | 1.250-1.270 |
| *R. pallescens* | 0.1467 | 0.9610 | 0.1524 | 1.017-1.077 |
| *R. neglectus* | 0.3099 | 0.8325 | 0.3431 | 1.151-1.174 |
| *R. nasutus* | 0.2346 | 0.8974 | 0.2569 | 1.103-1.154 |
| *R. ecuadoriensis* | 0.0855 | 0.6034 | 0.1061 | 1.187-1.358 |
| *R. colombiensis* | 0.0327 | 0.9293 | 0.0317 | 0.8547-1.101 |

\* Odds ratio between *Rhodnius* prevalence inside palm area (third column) and in the total area (first column).

Discriminating by palm species, *Rhodnius* species prevalence in areas with palms was much higher than in the entire area in some cases (Table 5). In *R. robustus*, prevalence increased more than eight times in areas with *As. aculeatum*, *As. murumuru* and *O. bataua* presence; in *R. prolixus*, it increased more than seven times in *Ac. aculeata* areas; in *R. pictipes*, more than six times in *As. aculeatum* and *At. maripa* areas; in *R. pallescens*, more than four times in *E. oleifera* areas; in *R. neglectus*, more than three times in *Sy. oleracea* areas; in *R. nasutus*, more than two times in *Cp. prunifera* areas; in *R. ecuadoriensis*, more than three times in *P. aequatorialis* areas; and in *R. colombiensis*, more than two times in *As. aculeatum* areas. In contrast, *Rhodnius* species prevalence was much lower in areas with certain palm species. For example, *R. prolixus* prevalence was very low in areas with *As. murumuru* and *R. pictipes* prevalence in areas with *Ac. aculeata* (Table 5). In this study, a *Rhodnius*-palm species pair was considered spatially associated if the odds ratio value was higher or equal to 2.

**Table 5.** *Rhodnius* species prevalence in different palm species areas.

| Species | *R. robustus* | *R. prolixus* | *R. pictipes* | *R. pallesc.* | *R. neglect.* | *R. nasutus* | *R. ecuador.* | *R. colomb.* |
| --- | --- | --- | --- | --- | --- | --- | --- | --- |
| *Ac. aculeat.* | 0.13<br>0.14-0.15 | **0.87**<br>**7.4-7.9** | 0.16<br>0.16-0.17 | 0.21<br>1.5-1.6 | **0.49**<br>**2.1-2.2** | 0.22<br>0.91-0.97 |  | 0.002<br>0.02-0.12 |
| *As. aculeat.* | **0.96**<br>**22.2-23.2** | 0.59<br>1.57-1.64 | **0.91**<br>**8.8-9.1** | 0.19<br>1.3-1.4 | 0.12<br>0.31-0.32 | 0.06<br>0.20-0.23 |  | **0.08**<br>**2.2-2.9** |
| *As. murum.* | **0.95**<br>**18.1-19.1** | 0.091<br>0.10-0.11 | **0.83**<br>**4.4-4.5** | 0.15<br>0.9-1.1 |  |  |  |  |
| *At. butyra.* | **0.82**<br>**4.3-4.4** | 0.33<br>0.54-0.56 | **0.76**<br>**2.8-2.9** | 0.18<br>1.2-1.3 |  |  | 0.002<br>0.01-0.03 | 0.05<br>1.3-1.8 |
| *At. maripa* | **0.93**<br>**12.1-12.5** | 0.35<br>0.58-0.60 | **0.88**<br>**6.4-6.5** | 0.18<br>1.3-1.4 | 0.13<br>0.33-0.34 | 0.15<br>0.56-0.62 | 0.002<br>0.01-0.03 | 0.02<br>0.64-0.91 |
| *At. phaler.* | 0.58<br>1.31-1.34 |  | 0.55<br>1.08-1.10 |  | **0.52**<br>**2.3-2.4** |  | 0.05<br>0.49-0.63 |  |
| *At. specio.* | 0.45<br>0.79-0.81 |  | 0.44<br>0.68-0.70 |  | **0.49**<br>**2.1-2.2** | 0.34<br>1.6-1.7 |  |  |
| *Cc. nucife.* | 0.54<br>1.11-1.13 | 0.54<br>1.26-1.30 | 0.54<br>1.02-1.04 | 0.15<br>1.0-1.1 | 0.27<br>0.8-0.9 | 0.32<br>1.5-1.6 | 0.14<br>1.7-1.9 | 0.03<br>0.86-1.11 |
| *Cp. prunif.* |  |  |  |  | 0.32<br>1.01-1.05 | **0.47**<br>**2.9-3.0** |  |  |
| *E. oleifer.* |  | 0.43<br>0.81-0.85 |  | **0.43**<br>**4.3-4.6** |  |  |  | 0.04<br>0.99-1.61 |
| *M. flexuos.* | **0.71**<br>**2.3-2.4** | 0.37<br>0.64-0.66 | 0.70<br>1.94-1.98 | 0.18<br>1.3-1.4 | 0.44<br>1.7-1.8 | 0.09<br>0.29-0.32 | 0.02<br>0.21-0.28 | 0.02<br>0.52-0.74 |
| *O. Bataua* | **0.89**<br>**8.1-8.4** | 0.27<br>0.40-0.41 | **0.82**<br>**3.9-4.0** | 0.18<br>1.2-1.3 |  |  | 0.04<br>0.45-0.55 | 0.01<br>0.22-0.36 |
| *P. aequat.* |  |  |  |  |  |  | **0.26**<br>**3.2-4.5** |  |
| *Sy. olerac.* | 0.001<br>0.001-0.002 |  |  |  | **0.61**<br>**3.4-3.5** |  |  |  |
| *Sy. orinoce.* |  |  |  | 0.07<br>0.38-0.45 |  |  |  | 0.001<br>0.02-0.09 |

Each cell: First line: *Rhodnius* species prevalence inside palm area. Second line: 95% confidence interval of the odds ratio between *Rhodnius* prevalence inside the palm area and in the total area. Only palm species with prevalence higher than 0.10 were included in the analysis. *Rhodnius*-palm pairs showing association (odds ratio higher than 2) are showing in bold.

To compare if *Rhodnius*-palm spatial association could be explained by niche similarity, niche overlap was compared between *Rhodnius*-palm pairs with and without spatial association (S2 Table). To this, n-dimensional hypervolumes overlapping was calculated by the function “dynRB_VPa” in the “dynRB” R package [86]. In all the *Rhodnius* species but *R. colombiensis*, mean niche overlap was higher in *Rhodnius*-palm pairs with spatial association than in pairs without the association.

Considering all *Rhodnius* and palm presence in the same extension (from Guatemala to northern Argentina), the highest *Rhodnius* richness (number of species) was concentrated in the Amazon region and the Guiana shield (Brazil, Colombia Venezuela and Guyana) (Fig 4). More than 60% of the area predicted for *Rhodnius* (i.e. area with at least one *Rhodnius* species), was predicted to be occupied by two or more *Rhodnius* species (Fig 4 up). In the limits of this region, only one *Rhodnius* species is predicted as present. The Amazon region was also the area with the highest predicted richness of palm species (species considered in this study), and 87% of the area predicted as present for palms (i.e. area with at least one palm species) is predicted to be occupied by two or more palm species. Almost all the considered area, from Guatemala to northern Argentina, had a continuous presence of palms (species with infestation reports).

**Fig. 4.**
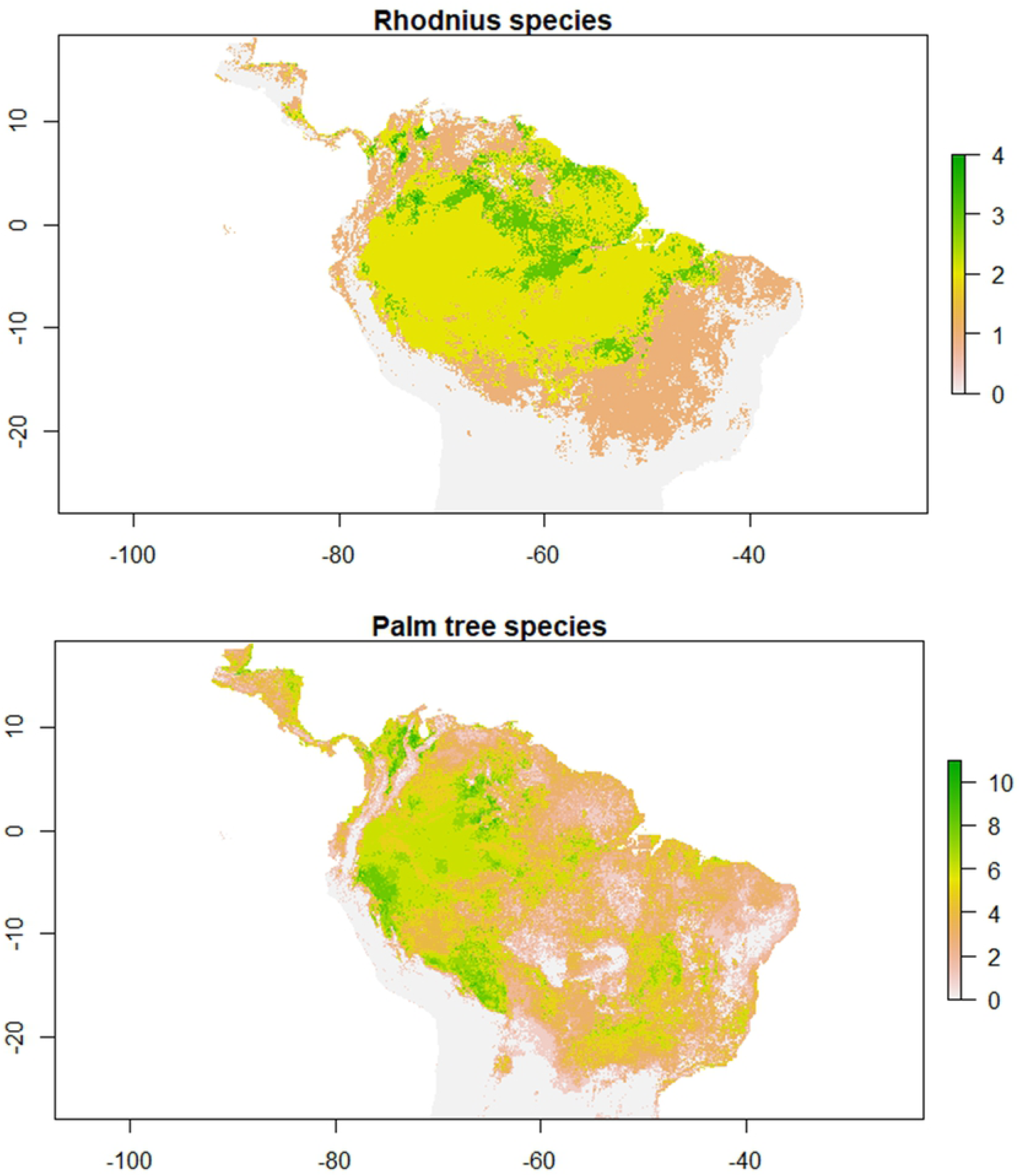
*Rhodnius* and palm tree species in the distribution area of the *Rhodnius* genus. Presence of each species were those predicted in at least three algorithms based on the 10% omission rate threshold. Horizontal axis: Longitude, Vertical axis: Latitude. Maps constructed with the raster R package [85].

Since both *Rhodnius* and palm predicted richness appear to be concentrated in the same region, the Amazon, spatial distribution was compared between *Rhodnius* and palms predicted richness. There is a likely relationship between *Rhodnius* and palms species richness (Fig. 5). Fifty percent of locations with four *Rhodnius* species had six to eight palm species. In contrast, fifty percent of locations with only one *Rhodnius* species had two to four palm species, and fifty percent of locations without *Rhodnius* species had no palm presence, or only one or two palm species.

**Fig 5.**
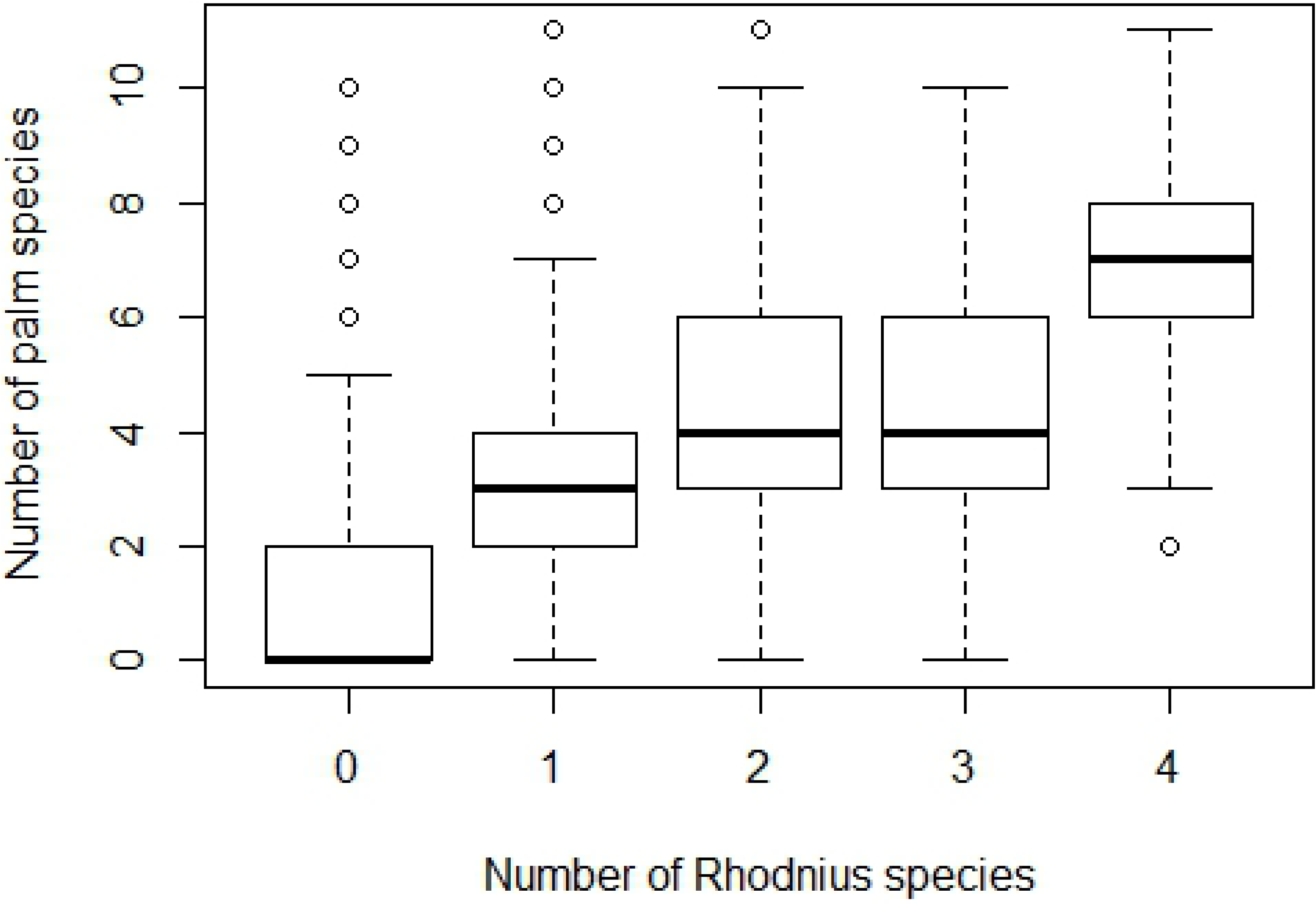
Spatial relationship between the number of *Rhodnius* and palm species.

### *Rhodnius* models with palm trees distributions as predictors

When *Rhodnius* models were run with palm distributions as predictors, performance behavior was similar to the previous models. Partial AUC ratios were significantly higher than the null model line except for *R. neglectus* and *R. colombiensis* (Table 2B); 10 percentile omission rates were higher than expected (and sometimes much higher), but zero percentile omission rates were closer to the expected values (Table 2B). The three species with the lowest occurrence number (*R. nasutus*, *R. colombiensis* and *R. ecuadoriensis*) had both omission rates very far from the expected values. As predictor, palm distributions showed to be not very relevant for *Rhodnius* models (Table 2B). Palm importance was low in all *Rhodnius* species but *R. pallescens* and *R. colombiensis*. In those species, however, the models did not show any increase in performance using palm distributions. Spatial differences in the predictions of models with and without palm distributions were scarce and disperse, and they are mainly located in the edges of the presence areas (Fig. 6).

**Fig. 6.**
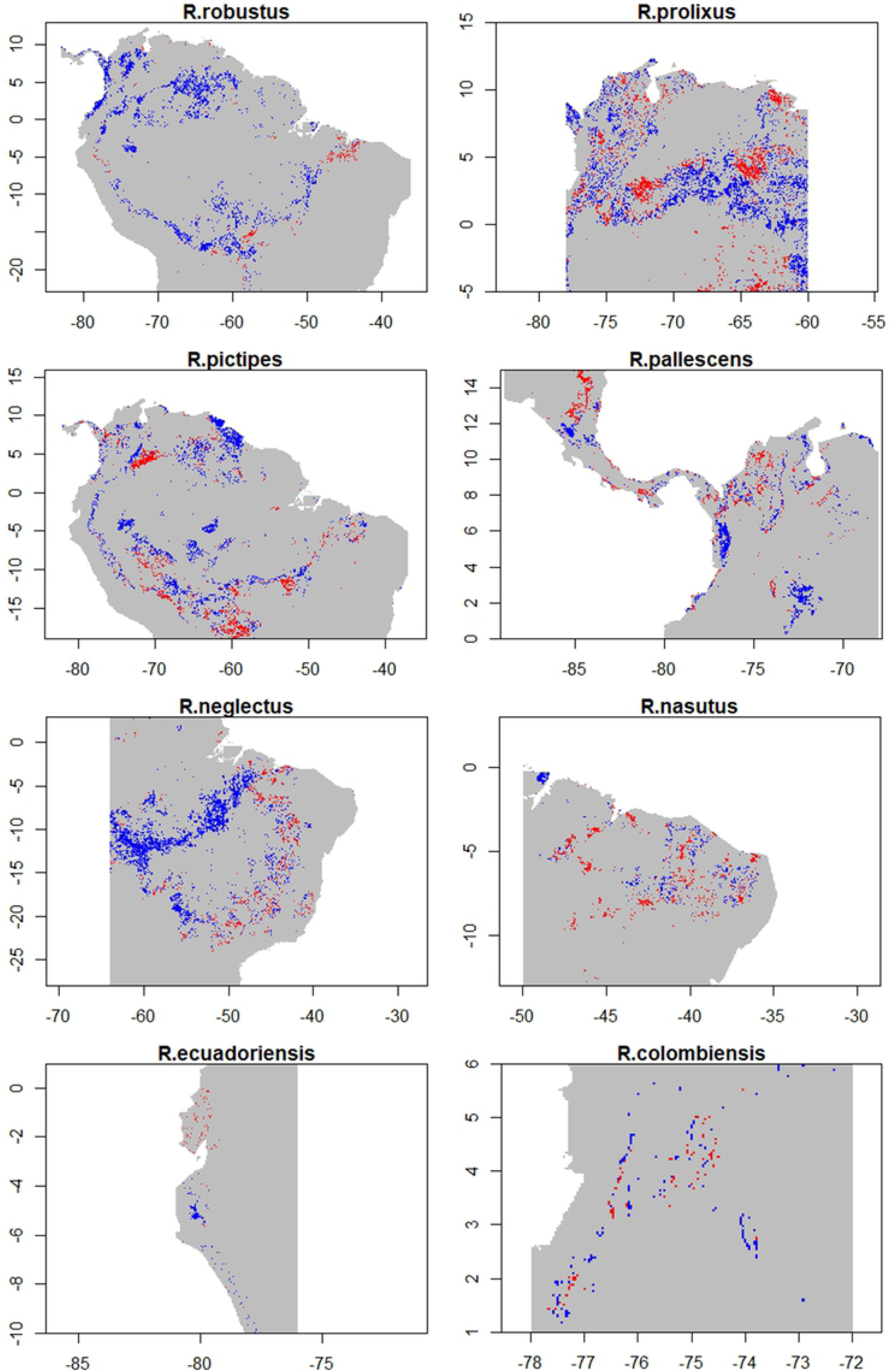
Comparison between *Rhodnius* models run with and without palm distributions as predictors. Red: Presences found only in models with palm distributions. Blue: Presences found only in models with environmental variables. Gray: Presences or absences predicted in both models. Presences were predicted at least three algorithms based on the 10% omission rate threshold. Horizontal axis: Longitude, Vertical axis: Latitude. Maps constructed with the raster R package [85].

## Discussion

Considering the association between *Rhodnius* species and palms, the prevalence of *Rhodnius* species was not much higher in palm tree areas than in the total modeled areas. That could be a consequence of the palm presence area, which was very big and, in some cases, encompassed all the extension. If palm area and total modeled area are similar, *Rhodnius* prevalence in both areas cannot be very different. That is demonstrated in the odd ratios values that were all very close to one. Therefore, based on *Rhodnius* prevalence comparison, the association between *Rhodnius* species and palm trees presence cannot be clearly determined.

In contrast, when palm species were considered, prevalence comparisons showed greater differences. Each *Rhodnius* species’ prevalence increased in specific palm areas compared to the entire area. Comparisons were several times higher in some cases (Table 5). That showed a clear spatial association between the presence of *Rhodnius* species and certain palm species. *Rhodnius* prevalence difference can be enormous between palm species. For instance, *R. robustus* presence was 150 times higher in *As. aculeatum* areas than in *Ac. aculeata* areas, and *R. prolixus* presence was 17 times higher in *A. aculeata* areas than in *O. bataua* areas. Hence, the palm species appears to be key for *Rhodnius*-palm association.

*Rhodnius*-palm spatial association could be explained by different causes: the ecological interaction between both organisms and the similarity in environmental factors that determine the distributions. Considering ecological interaction, palm species has demonstrated to be an important factor influencing the presence and abundance of triatomines in palm crowns [5,6,20,54,62,66]. Palm species differ in palm architecture, microclimatic conditions and the associated vertebrate fauna, factors that have been related to triatomine presence in palms [6]. Considering similarity in environmental factors, environmental variables as annual temperature, precipitation, and precipitation seasonality were important environmental factors for several *Rhodnius* and palm species distributions. Triatomines and palms have shown a high sensitivity to climatic conditions. Temperature affects physiological and behavioral processes of triatomines as egg production, hatching and immature development [87,88]. Climatic conditions could affect palm trees due to their soft and water-rich tissues, their inability to undergo dormancy and their general lack of mechanisms to avoid or tolerate frost [89]. Environmental similarity can be further to specific factors. Regarding the entire niche, overlap was higher in *Rhodnius*-palm pairs with spatial association than in those not associated. N-dimension volumes appeared to be more similar in certain *Rhodnius* and palm species, and that niche similarity can favor the co-occurrence. Both causes of Rhodnius and palm association, interaction and niche similarity, could be even complementary. Niche similarity between *Rhodnius* and palms could promote the presence of both organisms in the same location, and then, their ecological interaction would occur based on the advantages that palms offer to those hematophagous insects.

Analyzing by *Rhodnius* species*, R. robustus* distribution showed a great association with six palm species: *As. aculeatum*, *As. murumuru*, *At. butyracea*, *At. maripa*, *M. flexuosa* and *O. bataua*. The first four species have previous infestation reports by *R. robustus* [60,64– 67] while *M. flexuosa* and *O. bataua* don’t; however the presence of *R. robustus* is likely in these two palm species because they have an extended distribution in the Amazon region and reported infestations by other *Rhodnius* species. Even though *Ac. aculeata* has been found infested with *R. robustus* [64,65], spatial association was not found (Table 5). Those reports come from Venezuela where the six palm species showing association are absent. In that location, *R. robustus* bugs would associate with the palm species present in the region (i.e. *Ac. aculeata*).

*Rhodnius pictipes* results were very similar to those of *R. robustus,* displaying spatial association with the same palm species. That is caused by the similarity in both *Rhodnius* species distributions (Fig 1). Three species, *As. murumuru*, *At. maripa* and *O. bataua*, have previous infestation reports by *R. pictipes* [60]. The other palms, *As. aculeatum* and *At. butyracea*, have no reports but *R. pictipes* infestation can be considered as very possible based on the palm distributions and the reported infestation by other *Rhodnius* species [62,66,67].

Despite infesting several palm species (Table 1), *R. prolixus* only showed high spatial association with *Ac. aculeata*. No clear association was found with the other palms, and it could reflect the role of domiciliation in its distribution. In the *R. prolixus* occurrence set used in this study, several data could come from human dwellings, where bug populations can establish without the presence of close palms [90]. *Rhodnius prolixus* presence at high altitude locations (which have no palms) could be explained by the colonization of human dwellings [9]. Looking at *R. prolixus* predicted distribution its presence in highlands of Venezuela and Guiana could be considered as overprediction, since in that region, there are no occurrences nor predicted palm presence (Fig 1). However, those highlands are similar to the Andean zones where *R. prolixus* has been intensively reported.

Like *R. prolixus*, *R. pallescens* was also reported in several palm species (Table 1) but it only showed high association with *E. oleifera*. This palm has a broad distribution very similar to that of *R. pallescens* [19,36]. No association was found with *A. butyracea* even though this palm has several reports of *R. pallescens* infestation [52–55,57]. Lack of association with *A. butyracea* and other palm species could be caused by the presence of *R. pallescens* in other habitats such as armadillo burrows or by its presence in human dwellings [14] and chicken coops [58].

*Rhodnius nasutus* was also reported in several palm species (Table 1), but spatial association was found only with *Cp. prunifera* and *At. speciosa* (the latter with low odd ratio but close to 2). *Copernicia prunifera* was the most distributed palm species inside *R. nasutus* area, and their association have been frequently reported [5,49,91–94]. Considering the other palms with infestation reports, *Ac. aculeata* and *M. flexuosa* showed no association and *Sy. oleracea* prevalence was very small as to be considered in the analysis (lower than 0.10).

In contrast to the previous *Rhodnius* species, all the palm species associated with *R. neglectus* (*Ac. aculeata*, *At. phalerata*, *At. speciosa*, *M. flexuosa* and *Sy. oleracea*) were confirmed by previous infestation reports [6,51]. *Mauritia flexuosa* showed a lower odds ratio but closer to 2.

In *R. ecuadoriensis*, a high spatial association was seen with *P. aequatorialis*; this relationship has been deeply studied [47,48]. In the north of *R. ecuadoriensis* distribution, presence is related to palm trees presence (Fig. 3); while in the south, presence is related to domiciliation process with no palm trees [29,95]. In *R. colombiensis*, clear association was found with *As. aculeatum* and not with *At. butyracea*, the only species with infestation reports. Nevertheless, *R. colombiensis* distribution appears to be underestimated by the models, which produced very high omission rates (Table 2A).

Analyzing *Rhodnius* and palm distributions regionally, the Amazon appears to be a convergent area where several species intersected [28]. *Rhodnius*-palm association could be originated in the Amazon, and then, *Rhodnius* populations began a migration toward further zones in the north or the south progressing through zones with palm presence. From Mexico to Argentina, palm presence is continuous allowing the connection among zones. Further areas like the Andes and Central America in the north and Central Brazil in the South would have been colonized by *Rhodnius* bugs with an ongoing speciation process [96]. This agrees with the observation that only one *Rhodnius* species occurs in the limits of the complete *Rhodnius* genus distribution (Fig 4), while several species occur in the Amazon region [14].

High *Rhodnius* species richness shown in the Amazon appears to be related with high palm richness. Palms are considered to be suitable habitats for *Rhodnius* since they offer food and shelter. Habitat quality offered by palms could be heterogeneous among palm species [6], and ecological heterogeneity has been proposed as a driver of species richness for several reasons [97]: 1) Different habitat types could increase the available niche space and allow more species to coexist; 2) There would be more diversity in shelter and refuges from adverse environmental conditions and periods of climate change, promoting species persistence; 3) Speciation probability caused by isolation or adaptation to diverse environmental conditions should increase with greater ecological heterogeneity [97]. High diversity of palms, and therefore high diversity of habitats for *Rhodnius*, could favor the co-occurrence of several *Rhodnius* species in one specific region.

Palm distribution used as predictor did not increase model performance compared to the initial models. Palm presence information seems not to be relevant for *Rhodnius* models compared to environmental variables. In fact, the most important variables in models with and without palm distributions were usually similar. Species interactions are considered to affect species distribution mainly at local scales (e.g. landscape), and abiotic predictors, as temperature and precipitation, would affect species distributions in a bigger scale [98]. At the large spatial scale used in this study, palm distribution would not restrict *Rhodnius* distributions clearly; in a landscape scale however, results can be very different. The use of interacting species as predictors of *Rhodnius* distribution could need the inclusion of other participants like birds or mammal hosts. Those animals appear to be the crucial link between palms and *Rhodnius* triatomines [5]. Including a more complete scenario in the spatial modeling, mainly in the landscape scale, would increase the model complexity but could also increase predictability [98,99].

As distribution models, ENM are severely dependable on available information. The low occurrence number in some species and the biased distribution of information (e.g. some areas intensively sampled in comparison to others) could limit the validity of the conclusions. The presence of *Rhodnius* triatomines in more palm species than those considered in this study cannot be excluded, and the conclusions here are limited to one small subgroup of palm species inside the huge diversity of palm trees found in the *Rhodnius* presence area [19].

Considering model performance, evaluation showed a good discriminate power (pAUC) in most of the *Rhodnius* and palm species; however, in some cases, omission rates were higher than expected. This could be a consequence of the low occurrence number. To test models, 20% of the occurrences were used. In *R. colombiensis* for example, the size of the testing set was of four or five occurrences. If one or two points were omitted in one repetition, omission rates reach very high levels. That situation is less probable if occurrence number is big. With 372 occurrences, as in *R. prolixus*, more than 72 points are used for testing and leaving out two or three points do not increase omission levels dramatically.

High omission rates in the results have also shown to be related to a small species presence. Very small prevalence could be a consequence of model overfitting. Some algorithms, Random Forests for instance, gave models extremely fitted to the training data, with very high pAUC values but very low spatial transferability. On the contrary, MaxEnt models gave much lower omission rates, close to expected values, but suffer from high overprediction predicting presence in areas very different to the occurrences. Model averaging used in this study, which combined the outputs of five algorithms, could help to soften the performance limitations of each algorithm [83]. Comparing model performance, there was no algorithm being the best for all the species. For the same reason, agreement among algorithms was used in binary maps. The species presence was the coincidence of three or more algorithms instead of using only one algorithm.

To conclude, *Rhodnius* species showed to have a distributional association with certain palm species. Even though palms are widely distributed in every *Rhodnius* species range, the presence of a *Rhodnius* species is related to the presence of specific palm trees. Looking at a continental scale, this relationship could be linked to the *Rhodnius* origin, possibly in the Amazon region. Palm richness can be considered as an important factor allowing *Rhodnius* species co-occurrence. Despite the spatial association found, palm presence did not improve *Rhodnius* distribution models. The use of more interacting species as birds and mammal hosts could increase the complexity but also increase model performance and predictability.

## Acknowledgments

We thank Juan Manuel Cordovez (BIOMAC, Universidad de Los Andes, Colombia), Jorge Molina (CIMPAT, Universidad de Los Andes, Colombia) and Nicole L. Gottdenker (Department of Veterinary Pathology, University of Georgia, USA) for their helpful discussion and suggestions.

## Supporting information

**S1 Table. Performance statistics in *Rhodnius* ENM with the 16 first PCAs as environmental layers.** ^1^Median value for the five used algorithms. With each algorithm, value was the median of the ten repetitions carried out with different training and testing data set.

**S2 Table. Mean *Rhodnius*-palm niche overlap.** ^1^*Rhodnius*-palm species pair considered spatially associated if the odds ratio in table 5 was higher or equal to 2. Niche overlap ranges between 0 (No overlap) to 1 (complete overlap).

## References

1. WHO. Chagas disease (American trypanosomiasis) fact sheet (revised in June 2010). Wkly Epidemiol Rec. 2010;85: 334–336.

2. Gaunt M, Miles M. The Ecotopes and Evolution of Triatomine Bugs (Triatominae) and their Associated Trypanosomes. Mem Inst Oswaldo Cruz. 2000;95: 557–565. doi:10.1590/S0074-02762000000400019

3. Galvão C, Justi SA. An overview on the ecology of Triatominae (Hemiptera:Reduviidae). Acta Trop. Elsevier B.V.; 2015;151: 116–125. doi:10.1016/j.actatropica.2015.06.006

4. Lent H, Wygodzinsky P. Revision of the Triatominae (Hemiptera, Reduviidae), and Their Significance As Vectors of Chagas’ Diseas. Bull Am Museum Nat Hist. 1979;163: 123–520.

5. Dias FBS, Bezerra CM, De Menezes Machado EM, Casanova C, Diotaiuti L. Ecological aspects of Rhodnius nasutus Stål, 1859 (Hemiptera: Reduviidae: Triatominae) in palms of the Chapada do Araripe in Ceará, Brazil. Mem Inst Oswaldo Cruz. 2008;103: 824–830. doi:10.1590/S0074-02762008000800014

6. Abad-Franch F, Monteiro FA, Jaramillo O. N, Gurgel-Gonçalves R, Dias FBS, Diotaiuti L. Ecology, evolution, and the long-term surveillance of vector-borne Chagas disease: A multi-scale appraisal of the tribe Rhodniini (Triatominae). Acta Trop. 2009;110: 159–177. doi:10.1016/j.actatropica.2008.06.005

7. Teixeira ARL, Monteiro PS, Rebelo JM, Argañaraz ER, Vieira D, Lauria-Pires L, et al. Emerging chagas disease: Trophic network and cycle of transmission of Trypanosoma cruzi from palm trees in the Amazon. Emerg Infect Dis. 2001;7: 100– 112. doi:10.3201/eid0701.010115

8. Feliciangeli MD, Sanchez-Martin M, Marrero R, Davies C, Dujardin JP. Morphometric evidence for a possible role of Rhodnius prolixus from palm trees in house re-infestation in the State of Barinas (Venezuela). Acta Trop. 2007;101: 169– 177. doi:10.1016/j.actatropica.2006.12.010

9. Fitzpatrick S, Feliciangeli MD, Sanchez-Martin MJ, Monteiro FA, Miles MA. Molecular genetics reveal that silvatic Rhodnius prolixus do colonise rural houses. PLoS Negl Trop Dis. 2008;2: e210. doi:10.1371/journal.pntd.0000210

10. Guhl F, Pinto N, Marín D, Herrera C, Aguilera G, Naranjo JM, et al. Primer reporte de Rhodnius prolixus Stal, en Elaeis guineensis variedad Papúa, en plantaciones agroindustriales de Villanueva, Casanare. Biomedica. 2005;25: 158–159.

11. Carvalho DB, Almeida CE, Rocha CS, Gardim S, Mendonça VJ, Ribeiro AR, et al. A novel association between Rhodnius neglectus and the Livistona australis palm tree in an urban center foreshadowing the risk of Chagas disease transmission by vectorial invasions in Monte Alto City, São Paulo, Brazil. Acta Trop. 2014;130: 35– 38. doi:10.1016/j.actatropica.2013.10.009

12. Gamboa C. J. Comprobación de Rhodnius prolixus extradomiciliario en Venezuela. Bol la Of Sanit Panam. 1963; 18–25.

13. Sanchez-Martin MJ, Feliciangeli MD, Campbell-Lendrum D, Davies CR. Could the Chagas disease elimination programme in Venezuela be compromised by reinvasion of houses by sylvatic Rhodnius prolixus bug populations? Trop Med Int Heal. 2006;11: 1585–1593. doi:10.1111/j.1365-3156.2006.01717.x

14. Gorla D, Noireau F. Geographic distribution of Triatominae vectors in America. In: Telleria J, Tibayrenc M, editors. American Trypanosomiasis Chagas Disease One Hundred Years of Research. 2nd Editio. Elsevier; 2017. pp. 197–222.

15. Bargues MD, Schofield C, Dujardin JP. Classification and systematics of the Triatominae. In: Telleria J, Tibayrenc M, editors. American Trypanosomiasis Chagas Disease One Hundred Years of Research. Second edi. Amsterdam, Netherlands: Elsevier; 2017. pp. 113–144.

16. Carcavallo RU, Curto de Casas SI, Sherlock ÍA, Galíndez Girón I, Jurberg J, Galvao C, et al. Geographical distribution and alti-latitudinal dispersion. In: Carcavallo RU, Galíndez Girón I, Jurberg J, Lent H, editors. Atlas of Chagas disease in the Americas. Rio de Janeiro: Editora Fiocruz; 1999. pp. 747–792.

17. Galvao C, Carcavallo R, Da Silva Rocha D, Jurberg J. A checklist of the current valid species of the subfamiliy Triatominae Jeannel, 1919 (Hemiptera, Reduviidae) and their geographical distribution, with nomeclatural and taxonomic notes. Zootaxa. 2003;201: 1–36.

18. Galeano G, Bernal R. Palmas de Colombia. Guía de campo. Bogotá D.C.: Editorial Universidad Nacional de Colombia; 2010.

19. Henderson A, Galeano G, Bernal R. Field Guide to the Palm of the Americas. New Jersey, USA: Princeton University Press; 1995.

20. Abad-Franch F, Lima MM, Sarquis O, Gurgel-Gonçalves R, Sánchez-Martín M, Calzada J, et al. On palms, bugs, and Chagas disease in the Americas. Acta Trop. 2015;151: 126–141. doi:10.1016/j.actatropica.2015.07.005

21. Urbano P, Poveda C, Molina J. Effect of the physiognomy of Attalea butyracea (Arecoideae) on population density and age distribution of Rhodnius prolixus (Triatominae). Parasit Vectors. 2015;8: 1–12. doi:10.1186/s13071-015-0813-6

22. Angulo VM, Esteban L, Luna KP. Attalea butyracea próximas a las viviendas como posible fuente de infestación domiciliaria por Rhodnius prolixus (Hemiptera: Reduviidae) en los Llanos Orientales de Colombia. Biomédica. 2012;32: 277–285. doi:10.7705/biomedica.v32i2.430

23. Gottdenker NL, Streicker DG, Faust CL, Carroll CR. Anthropogenic Land Use Change and Infectious Diseases: A Review of the Evidence. Ecohealth. 2014; 619– 632. doi:10.1007/s10393-014-0941-z

24. Galindez I, Curto de Casas SI, Carcavallo RU, Jurberg J, Mena Segura CA. Geographical distribution and alti-latitudinal dispersion of the tribe Rhodniini (Hemiptera, Reduviidae, Triatominae). Entomol y Vectores. 1996;3: 3–20.

25. Molina J a, Gualdrón LE, Brochero HL, Olano V, Barrios D, Guhl F, et al. Distribución actual e importancia epidemiológica de las especies de triatominos (Reduviidae: Triatominae) en Colombia. Biomédica. 2000;20: 344–360. doi:10.7705/biomedica.v20i4.1078

26. Guhl F, Aguilera G, Pinto N, Vergara D. Actualización de la distribución geográfica y ecoepidemiología de la fauna de triatominos (Reduviidae: Triatominae) en Colombia. Biomédica. 2007;27: 143–162. doi:http://dx.doi.org/10.7705/biomedica.v27i1.258

27. Cuba Cuba CA, Abad-Franch F, Rodríguez JR, Vásquez FV, Velasquez LP, Miles MA. The triatomines of Northern Peru, with emphasis on the ecology and infection by trypanosomes of Rhodnius ecuadoriensis (Triatominae). Mem Inst Oswaldo Cruz. 2002;97: 175–183. doi:10.1590/S0074-02762002000200005

28. Abad-Franch F, Monteiro FA. Biogeography and Evolution of Amazonian Triatomines (Heteroptera: Reduviidae): Implications for Chagas Disease Surveillance in Humid Forest Ecoregions. Mem Inst Oswaldo Cruz. 2007;102: 57– 69.

29. Abad-Franch F, Paucar C A, Carpio C C, Cuba Cuba CA, Aguilar V HM, Miles MA. Biogeography of triatominae (Hemiptera: Reduviidae) in Ecuador: Implications for the design of control strategies. Mem Inst Oswaldo Cruz. 2001;96: 611–620. doi:10.1590/S0074-02762001000500004

30. Lomolino M V., Riddle BR, Whittaker RJ, Brown JH. Biogeography. Fourth Edi. Sinauer Associates, Inc.; 2010.

31. Franklin J. Mapping Species Distribution. Cambridge, UK: Cambridge University Press; 2009.

32. Batista TA, Gurgel-Goncalves R. Ecological niche modelling and differentiation between Rhodnius neglectus Lent, 1954 and Rhodnius nasutus Stal, 1859 (Hemiptera: Reduviidae: Triatominae) in Brazil. Mem Inst Oswaldo Cruz. 2009;104: 1165–1170. doi:10.1590/S0074-02762009000800014

33. Gurgel-Goncalves R, Cuba CAC. Predicting the potential geographical distribution of Rhodnius neglectus (Hemiptera, Reduviidae) based on ecological niche modeling. J Med Entomol. 2009;46: 952–60. doi:10.1603/033.046.0430

34. Gurgel-Gonçalves R, Galvão C, Costa J, Peterson AT. Geographic distribution of chagas disease vectors in Brazil based on ecological niche modeling. J Trop Med. 2012;2012: 1–15. doi:10.1155/2012/705326

35. Pereira JM, de Almeida PS, de Sousa AV, de Paula AM, Bomfim Machado R, Gurgel-Gonçalves R. Climatic factors influencing triatomine occurrence in Central-West Brazil. Mem Inst Oswaldo Cruz. 2013;108: 335–341. doi:10.1590/0074-0276108032013012

36. Arboleda S, Gorla DE, Porcasi X, Saldaña A, Calzada J, Jaramillo-O N. Development of a geographical distribution model of Rhodnius pallescens Barber, 1932 using environmental data recorded by remote sensing. Infect Genet Evol. 2009;9: 441–448. doi:10.1016/j.meegid.2008.12.006

37. Parra-Henao G, Suárez-Escudero LC, González-Caro S. Potential Distribution of Chagas Disease Vectors (Hemiptera, Reduviidae, Triatominae) in Colombia, Based on Ecological Niche Modeling. J Trop Med. 2016;2016: 1–10. doi:10.1155/2016/1439090

38. Bustamante DM, Monroy MC, Rodas AG, Juarez JA, Malone JB. Environmental determinants of the distribution of Chagas disease vectors in south-eastern Guatemala. Geospat Health. 2007;1: 199–211. doi:10.4081/gh.2007.268

39. Araújo MB, Luoto M. The importance of biotic interactions for modelling species distributions under climate change. Glob Ecol Biogeogr. 2007;16: 743–753. doi:10.1111/j.1466-8238.2007.00359.x

40. Heikkinen RK, Luoto M, Virkkala R, Pearson RG, Körber JH. Biotic interactions improve prediction of boreal bird distributions at macro-scales. Glob Ecol Biogeogr. 2007;16: 754–763. doi:10.1111/j.1466-8238.2007.00345.x

41. Meier ES, Kienast F, Pearman PB, Svenning JC, Thuiller W, Araújo MB, et al. Biotic and abiotic variables show little redundancy in explaining tree species distributions. Ecography (Cop). 2010;33: 1038–1048. doi:10.1111/j.1600-0587.2010.06229.x

42. Schweiger O, Heikkinen RK, Harpke A, Hickler T, Klotz S, Kudrna O, et al. Increasing range mismatching of interacting species under global change is related to their ecological characteristics. Glob Ecol Biogeogr. 2012;21: 88–99. doi:10.1111/j.1466-8238.2010.00607.x

43. Leathwick JR, Austin MP. Competitive interactions between tree species in New Zealand’s old-growth indigenous forests. Ecology. 2001;82: 2560–2573. doi:10.1890/0012-9658(2001)082[2560:CIBTSI]2.0.CO;2

44. Leathwick JR. Intra-generic competition among Nothofagus in New Zealand’s primary indigenous forests. Biodivers Conserv. 2002;11: 2177–2187. doi:10.1023/A:1021394628607

45. Pellissier L, Anne Bråthen K, Pottier J, Randin CF, Vittoz P, Dubuis A, et al. Species distribution models reveal apparent competitive and facilitative effects of a dominant species on the distribution of tundra plants. Ecography (Cop). 2010;33: 1004–1014. doi:10.1111/j.1600-0587.2010.06386.x

46. Arévalo A, Carranza JC, Guhl F, Clavijo JA, Vallejo GA. Comparación del ciclo de vida de Rhodnius colombiensis Moreno, Jurberg & Galvão, 1999 y Rhodnius prolixus Stal, 1872 (Hemiptera, Reduviidae, Triatominae) en condiciones de laboratorio. Biomédica. 2007;27: 119–29.

47. Noireau F, Abad-Franch F, Valente SAS, Dias-Lima A, Lopes CM, Cunha V, et al. Trapping Triatominae in Silvatic Habitats. Memias Do Inst Oswaldo Cruz. 2002;97: 61–63. doi:10.1590/S0074-02762002000100009

48. Abad-Franch F, Palomeque FS, Aguilar VHM, Miles MA. Field ecology of sylvatic Rhodnius populations (Heteroptera, Triatominae): risk factors for palm tree infestation in western Ecuador. Trop Med Int Heal. 2005;10: 1258–1266. doi:10.1111/j.1365-3156.2005.01511.x

49. Dias FBS, de Paula AS, Belisário CJ, Lorenzo MG, Bezerra CM, Harry M, et al. Influence of the palm tree species on the variability of Rhodnius nasutus Stål, 1859 (Hemiptera, Reduviidae, Triatominae). Infect Genet Evol. 2011;11: 869–877. doi:10.1016/j.meegid.2011.02.008

50. Gurgel-Gonçalves R, Cuba C a. C. Estrutura de populações de Rhodnius neglectus Lent e Psammolestes tertius Lent & Jurberg (Hemiptera, Reduviidae) em ninhos de pássaros (Furnariidae) presentes na palmeira Mauritia flexuosa no Distrito Federal, Brasil. Rev Bras Zool. 2007;24: 157–163. doi:10.1590/S0101-81752007000100019

51. Gurgel-Gonçalves R, Duarte MA, Ramalho ED, Torre Palma AR, Romaña CA, Cuba-cuba CA. Distribuição espacial de populações de triatomíneos (Hemiptera: Reduviidae) em palmeiras da espécie Mauritia flexuosa no Distrito Federal, Brasil. Rev Soc Bras Med Trop. 2004;37: 241–247. doi:10.1590/S0037-86822004000300010

52. Whitlaw JT, Chaniotis BN. Palm trees and Chagas’ disease in Panama. Am J Trop Med Hyg. 1978;27: 873–881. doi:10.4269/ajtmh.1978.27.873

53. Pizarro Novoa JC, Romaña C. Variación estacional de una población silvestre de Rhodnius pallescens Barber 1932 (Heteroptera: Triatomiane) en la costa caribe colombiana. Bull Inst Fr Études Andin. 1998;27: 309–325.

54. Romaña CA, Pizarro JC, Rodas E, Guilbert E. Palm trees as ecological indicators of risk areas for Chagas disease. Trans R Soc Trop Med Hyg. 1999;93: 594–595. doi:10.1016/S0035-9203(99)90059-7

55. Salazar DA, Calle J. Caracterización ecoepidemiológica de Rhodnius pallescens en la palma Attalea butyracea en la región Momposina (Colombia). Actual Biológicas. 2003;25: 31–38.

56. Gottdenker NL, Chaves LF, Calzada JE, Salda??a A, Carroll CR. Host Life History Strategy, Species Diversity, and Habitat Influence Trypanosoma cruzi Vector Infection in Changing Landscapes. PLoS Negl Trop Dis. 2012;6: 5–7. doi:10.1371/journal.pntd.0001884

57. Cantillo-Barraza O, Chaverra D, Marcet P, Arboleda-Sánchez S, Triana-Chávez O. Trypanosoma cruzi transmission in a Colombian Caribbean region suggests that secondary vectors play an important epidemiological role. Parasit Vectors. 2014;7: 381. doi:10.1186/1756-3305-7-381

58. Jaramillo N, Schofield CJ, Gorla D, Caro-Riaño H, Moreno J, Mejia E, et al. The Role of Rhodnius Pallescens as a Vector of Chagas Disease in Colombia and Panama. Res Rev Parasitol. 2000;60: 75–82.

59. Gómez-Núñez JC. Resting places, dispersal and survival of CO60-tagged adult Rhodnius prolixus. J Med Entomol. 1969;6: 83–86.

60. Ricardo-Silva AH, Lopes CM, Ramos LB, Marques WA, Mello CB, Duarte R, et al. Correlation between populations of Rhodnius and presence of palm trees as risk factors for the emergence of Chagas disease in Amazon region, Brazil. Acta Trop. 2012;123: 217–223. doi:10.1016/j.actatropica.2012.05.008

61. Fitzpatrick SO. The analysis of the relationship between domestic and silvatic populations of Rhodnius prolixus(Hemiptera : Reduviidae) in Venezuela by geometric morphometric and molecular methods. London School of Hygiene & Tropical Medicine. 2007. doi:10.17037/PUBS.00682357

62. D’Alessandro A, Barreto P, Saravia N, Barreto M. Epidemiology of Trypanosoma cruzi in the Oriental Plains of Colombia. Am J Trop Med Hyg. 1984;33: 1084–1095.

63. Morocoima A, Chique J, Zavala-Jaspe R, Díaz-Bello Z, Ferrer E, Urdaneta-Morales S, et al. Commercial coconut palm as an ecotope of Chagas disease vectors in north-eastern Venezuela. J Vector Borne Dis. 2010;47: 76–84.

64. Longa ANA, Scorza V. Acrocomia aculeata (Palmae), hábitat silvestre de Rhodnius robustus en el Estado Trujillo, Venezuela. Parasitol Latinoam. 2005;60: 17–24.

65. Longa A, Scorza JV. Migración de Rhodnius robustus (Hemiptera: Triatominae) desde Acrocomia aculeata (Palmae) hacia domicilios rurales en Venezuela. Boletín Malariol y Salud Ambient. 2007;47: 213–220.

66. Dias FBS, Quartier M, Diotaiuti L, Mejía G, Harry M, Lima ACL, et al. Ecology of Rhodnius robustus Larrousse, 1927 (Hemiptera, Reduviidae, Triatominae) in Attalea palm trees of the Tapajós River Region (Pará State, Brazilian Amazon). Parasit Vectors. 2014;7: 154. doi:10.1186/1756-3305-7-154

67. Feliciangeli MD, Dujardin JP, Bastrenta B, Mazzarri M, Villegas J, Flores M, et al. Is Rhodnius robustus (hemiptera: Reduviidae) responsible for chagas disease transmission in western venezuela? Trop Med Int Heal. 2002;7: 280–287. doi:10.1046/j.1365-3156.2002.00853.x

68. Ceccarelli S, Balsalobre A, Medone P, Cano ME, Gonçalves RG, Feliciangeli D, et al. Data Descriptor: DataTri, a database of American triatomine species occurrence. Sci Data. 2018;5: 1–9. doi:10.1038/sdata.2018.71

69. Hijmans RJ, Phillips S, Leathwick J, Elith J. dismo: Species Distribution Modeling. R package version 1.1–4. 2017.

70. Telleria J, Tibayrenc M, editors. American Trypanosomiasis Chagas Disease. One Hundred Years of Research. Second. Amsterdam, Netherlands: Elsevier; 2017.

71. Chávez J. Contribución al estudio de los triatominos del Perú: Distribución geográfica, nomenclatura y notas taxonómicas. An la Fac Med. 2006;67: 65–76. doi:http://dx.doi.org/10.15381/anales.v67i1.1296

72. Sarquis O, Carvalho-Costa FA, Toma HK, Georg I, Burgoa MR, Lima MM. Eco-epidemiology of Chagas disease in northeastern Brazil: Triatoma brasiliensis, T. pseudomaculata and Rhodnius nasutus in the sylvatic, peridomestic and domestic environments. Parasitol Res. 2012;110: 1481–1485. doi:10.1007/s00436-011-2651-6

73. Moraes R. M. Ecología y formas de vida de las palmas bolivianas. Ecol en Boliv. 1989;13: 33–45.

74. Guhl F. Geographical distribution of Chagas disease. In: Telleria J, Tibayrenc M, editors. American Trypanosomiasis Chagas Disease One Hundred Years of Research. Second. Elsevier; 2017. pp. 89–112.

75. Hashimoto K, Schofield CJ. Elimination of Rhodnius prolixus in Central America. Parasite. 2012;5: 45. doi:10.1186/PREACCEPT-1790853046685273

76. Aiello-Lammens ME, Boria RA, Radosavljevic A, Vilela B, Anderson RP. spThin: An R package for spatial thinning of species occurrence records for use in ecological niche models. Ecography (Cop). 2015;38: 541–545. doi:10.1111/ecog.01132

77. Hijmans RJ, Cameron SE, Parra JL, Jones PG, Jarvis A. Very high resolution interpolated climate surfaces for global land areas. Int J Climatol. 2005;25: 1965– 1978. doi:10.1002/joc.1276

78. Hastings DA, Dunbar PK, Elphingstone, Gerald M. Bootz M, Murakami H, Maruyama H, Masaharu H, et al. The Global Land One-kilometer Base Elevation (GLOBE) Digital Elevation Model, Version 1.0. Boulder, Colorado, U.S.A.: National Oceanic and Atmospheric Administration, National Geophysical Data Center; 1999.

79. Hay SI, Tatem AJ, Graham AJ, Goetz SJ, Rogers DJ. Global Environmental Data for Mapping Infectious Disease Distribution. Adv Parasitol. 2006;62: 37–77. doi:10.1016/S0065-308X(05)62002-7

80. Thuiller W, Georges D, Engler R, Breiner F. biomod2: Ensemble Platform for Species Distribution Modeling. R package version 3.3–7. 2016.

81. Warren DL, Seifert S. Ecological niche modeling in Maxent: the importance of model complexity and the performance of model selection criteria. Ecol Soc Am. 2011;21: 335–342. doi:10.1890/10-1171.1

82. Muscarella R, Galante PJ, Soley-Guardia M, Boria RA, Kass JM, Uriarte M, et al. ENMeval: An R package for conducting spatially independent evaluations and estimating optimal model complexity for Maxent ecological niche models. Methods Ecol Evol. 2014;5: 1198–1205. doi:10.1111/2041-210X.12261

83. Guisan A, Thuiller W, Zimmermann NE. Habitat Suitability and Distribution Models. With Applications in R. Cambridge University Press; 2017.

84. Peterson AT, Papeş M, Soberón J. Rethinking receiver operating characteristic analysis applications in ecological niche modeling. Ecol Modell. 2008;213: 63–72. doi:10.1016/j.ecolmodel.2007.11.008

85. Hijmans RJ. raster: Geographic Data Analysis and Modeling. R package version 2.6–7. 2017.

86. Schreyer M, Trutschnig W, Junker RR, Kuppler J, Bathke A, Parkinson JH, et al. dynRB: Dynamic Range Boxes. 2018.

87. Luz C, Fargues J, Grunewald J. Development of Rhodnius prolixus (Hemiptera: Reduviidae) under Constant and Cyclic Conditions of Temperature and Humidity. Mem Inst Oswaldo Cruz. 1999;94: 403–409. doi:10.1590/S0074-02761999000300022

88. Clark N. The Effect of Temperature and Humidity upon the Eggs of the Bug, Rhodnius prolixus (Heteroptera, Reduviidae). J Anim Ecol. 1935;4: 82–87.

89. Tomlinson PB. The uniqueness of palms. Bot J Linn Soc. 2006;151: 5–14. doi:10.1111/j.1095-8339.2006.00520.x

90. Catalá SS, Noireau F, Dujardin JP. Biology of Triatominae. In: Telleria J, Tibayrenc M, editors. American Trypanosomiasis Chagas Disease One Hundred Years of Research. Second Edi. Amsterdam, Netherlands: Elsevier; 2017. pp. 145–168.

91. Lima MM, Coutinho CFS, Gomes TF, Oliveira TG, Duarte R, Borges-Pereira J, et al. Risk presented by Copernicia prunifera palm trees in the Rhodnius nasutus distribution in a Chagas disease-endemic area of the Brazilian northeast. Am J Trop Med Hyg. 2008;79: 750–754.

92. Lima MM, Sarquis O. Is Rhodnius nasutus (Hemiptera; Reduviidae) changing its habitat as a consequence of human activity? Parasitol Res. 2008;102: 797–800. doi:10.1007/s00436-007-0823-1

93. Lima MM, Sarquis O, Oliveira TG de, Gomes TF, Coutinho C, Daflon-Teixeira NF, et al. Investigation of Chagas disease in four periurban areas in northeastern Brazil: Epidemiologic survey in man, vectors, non-human hosts and reservoirs. Trans R Soc Trop Med Hyg. 2012;106: 143–149. doi:10.1016/j.trstmh.2011.10.013

94. Sarquis O, Borges-Pereira J, Mac Cord JR, Ferreira Gomes T, Cabello PH, Lima MM. Epidemiology of Chagas Disease in Jaguaruana, Ceará, Brazil. I. Presence of Triatomines and Index of Trypanosoma cruzi Infection in Four Localities of a Rural Area. Memorias do Inst Oswaldo Cruz Rio Janeiro. 2004;99: 263–270. doi:10.1097/NNE.0000000000000486

95. Vargas F, Córdova Paz Soldán O, Marín C, Jose Rosales M, Sánchez-Gutierrez R, Sánchez-Moreno M. Epidemiology of American trypanosomiasis in northern Peru. Ann Trop Med Parasitol. 2007;101: 643–648. doi:10.1179/136485907X229031

96. Schofield CJ, Dujardin JP. Theories on the evolution of Rhodnius. Actual Biol. 1999;21: 183–197.

97. Stein A, Gerstner K, Kreft H. Environmental heterogeneity as a universal driver of species richness across taxa, biomes and spatial scales. Ecol Lett. 2014;17: 866–880. doi:10.1111/ele.12277

98. Peterson AT. Mapping disease transmission risk. Baltimore, USA: Johns Hopkins University Press; 2014.

99. Samy AM, van de Sande WWJ, Fahal AH, Peterson AT. Mapping the Potential Risk of Mycetoma Infection in Sudan and South Sudan Using Ecological Niche Modeling. PLoS Negl Trop Dis. 2014;8. doi:10.1371/journal.pntd.0003250

